# A contraction-reaction-diffusion model for circular pattern formation in embryogenesis

**DOI:** 10.1101/2021.05.14.444097

**Authors:** Tiankai Zhao, Yubing Sun, Xin Li, Mehdi Baghaee, Yuenan Wang, Hongyan Yuan

## Abstract

Reaction-diffusion models have been widely used to elucidate pattern formation in developmental biology. More recently, they have also been applied in modeling cell fate patterning that mimic early-stage human development events utilizing geometrically confined pluripotent stem cells. However, the traditional reaction-diffusion equations could not satisfactorily explain the concentric ring distributions of various cell types, as they do not yield circular patterns even for circular domains. In previous mathematical models that yield ring patterns, certain conditions that lack biophysical understandings had been considered in the reaction-diffusion models. Here we hypothesize that the circular patterns are the results of the coupling of the mechanobiological factors with the traditional reaction-diffusion model. We propose two types of coupling scenarios: tissue tension-dependent diffusion flux and traction stress-dependent activation of signaling molecules. By coupling reaction-diffusion equations with the elasticity equations, we demonstrate computationally that the contraction-reaction-diffusion model can naturally yield the circular patterns.

## Introduction

The reaction-diffusion models have been widely used to explain pattern formation in developmental biology, where an activator-inhibitor network dominates the certain stage in biological development and leads to Turing Patterns afterwards. For example, Kondo et al^1^applied the model to study the movement of zebrafish strips in a recovery process. More recently, Human pluripotent stem cells (hPSCs)-based in vitro models have been developed to study early-stage human development^2–7^ (see an experimental result in Fig. 1). In these models, hPSCs were confined in mesoscale circular patterns and exposed to uniform morphogens or small molecules in culture media. Under these conditions, hPSCs differentiated into self-organized concentric rings of different cell types, mimicking the formation of three germ layers during gastrulation, or neuroepithelium/neural crest/epidermis patterning during neural plate induction. To explain such autonomous cell patterning behaviors, reaction-diffusion models have also been applied in these in vitro systems. Tewary et al^3^ studied the self-organized human peri-gastrulation-like-pattering, and suggested that the three germ layers formed in confined hPSCs were induced by reaction-diffusion of BMP4 and its inhibitor Noggin, by using a stepwise reaction-diffusion model. We recently reported that the patterning of neuroepithelium, neural crest, and epidermis can be tuned by temporally modulating the reaction-diffusion of BMP-Noggin and Wnt-Dkk pairs^6^.

**Figure 1.**
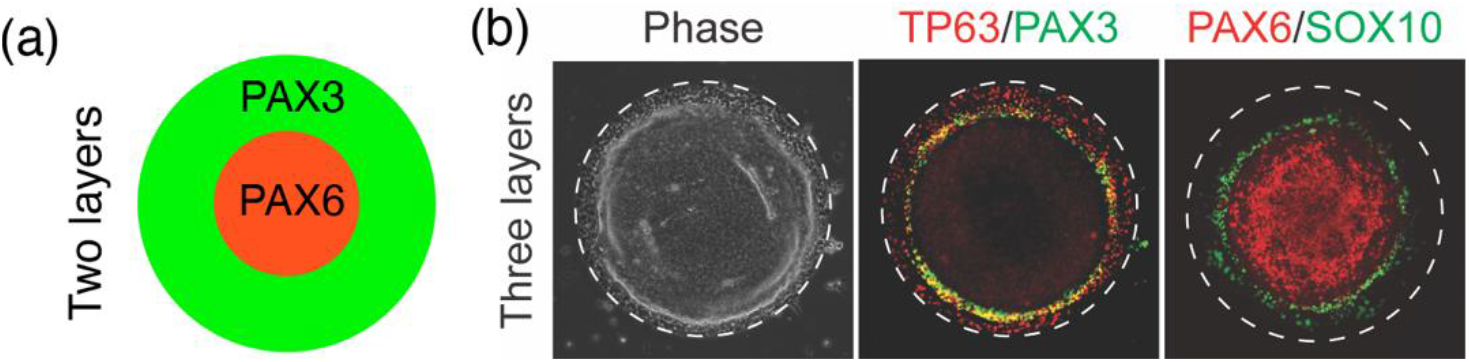
Schematic illustration and fluorescence images showing the self-organized ring-structure of neuroepithelium, neural crest and epidermis formed on micropatterned substrate. (a) The schematic illustration of a two-layer structure of the work by Xue et al.^4^; (b) A three-layer-structure showing the epidermis markers (TP63), the neuroepithelial cell marker (PAX6), and the neural crest markers (SOX10, PAX3), (Reprinted with permission from Xie et al.^6^).

The basic reaction-diffusion models do not predict axisymmetric patterns in a circular domain, instead they only produce dotted or stripped patterns, as shown in Fig. 2. While reaction-diffusion equations were generally successful in explaining the orders of different cell types in circular patterns, to produce circular patterns observed in experiments, previous studies have incorporated additional mathematical treatments that lack biophysical relevance into reaction-diffusion models. Namely, In Sanderson et al.’s work^8^, axisymmetric diffusion constant or reaction rate coefficient were directly defined. Tewary et al^3^. imposed circular initial conditions without direct experimental evidence, and Xie et al^6^. computationally modeled the pattern generation using an axisymmetric model directly. Therefore, the physical mechanism for the robust formation of concentric rings of different cell types across various models is still poorly understood.

**Figure 2.**
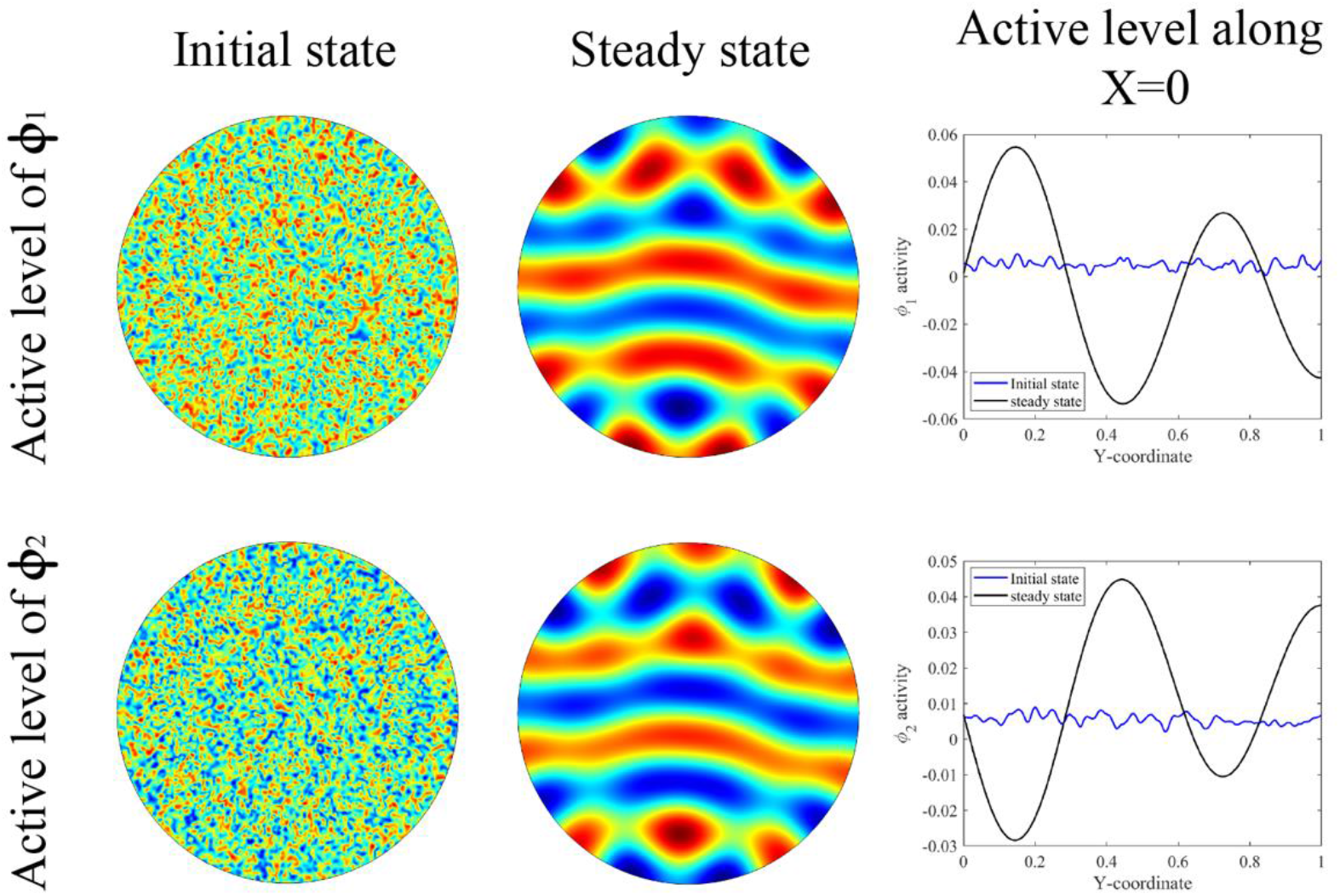
The non-axisymmetric Turing pattern contour generated by the basic reaction-diffusion model (red for large, and blue for small) and the active levels of the activators and inhibitors along the Y-axis at the initial and steady states.

A number of previous works have studied the cellular mechanobiology^9–11^, such as stress-dependent pattern formation^12–16^. Nelson et al^17^ have shown a correspondence between the patterns of proliferative foci regions and the areas of high traction forces. Yuan et al studied the maximum principal-stress-guided myofibril organizations^14^ and developed a contraction-reaction-diffusion model for morphological study of single-cell migration^15^. In these works, the solution of the elasticity equation for a contractile tissue, in general, predict a tension distribution highest in the middle of the domain and vanish at the free edge, and a traction force distribution highest at the edge and decreases towards the center. We hypothesize that this radial distribution feature can naturally lead to circular/ring pattern.

Here we propose that circular patterns are caused by the combination of mechanical stresses presented in the cellular tissue and the reaction-diffusion processes of signaling molecules. Specifically, we hypothesize two possible mechanisms that are (1) the tissue-tension-driven diffusion of signaling molecules and (2) the traction-forces-mediated activation of signaling molecules. By coupling reaction-diffusion equations with elasticity equations, we demonstrate computationally that the contraction-reaction-diffusion model can naturally yield circular patterns.

### Diffusion-driven Turing Instability for signaling mechanism in biological tissue development

The pattern formations in the process of tissue development, such as skin patterns of zebrafish^1^ and limbs development^18,19^, have been described by reaction-diffusion models in the literature^20^:

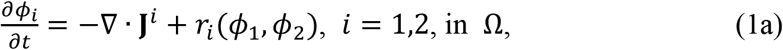

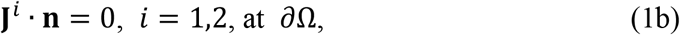

where

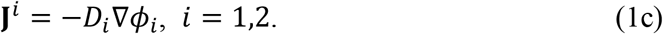

The active levels of the activators and inhibitors: two signaling molecules generated in the process of the tissue development, are described by the concentration of the two variables *ϕ*_1_ and *ϕ*_2_ in the diffusion-reaction equation. The parameters *D*_*i*_ (*i* = 1,2) are the overall diffusivity with respect to the active levels of the two kinds of the molecules. Their values are set in the following way: *D*_1_ = *hD*, *D*_2_ = *D*, where *D* and *h* are model parameters. The nonlinear reaction term is expanded in a Taylor’s series to third orders with some terms neglected:

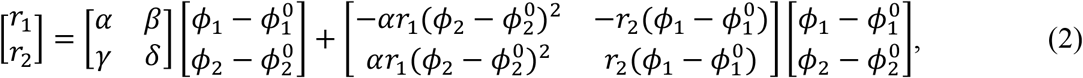

Where *α*, *β*, *γ*, *δ*, *r*_1_, and *r*_2_ are model parameters. The initial values of the two components (*ϕ*_1_, *ϕ*_2_) are set to be 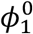 and 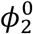, respectively.

For the basic reaction-diffusion model defined in Eq. (1)–(2), the following conditions must be satisfied for Turing instability to occur^20^:

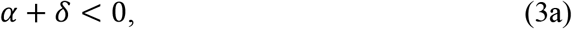

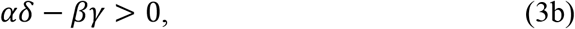

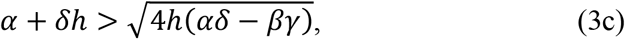

The wavenumber-related parameter at which the instability grows fastest is approximately given as:

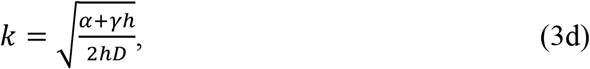

All the parameters are in the dimensionless form. The equations are all solved by the COMSOL Multiphysics by finite elements method, and an adaptive time integration scheme is used for the time-dependent problem. The quadratic triangular elements are adopted to generate the fine mesh for our problem.

Figure 2 shows the simulation results of the reaction-diffusion model (i.e., Eq. (1) and (2)). The simulation is carried out in a circular domain with a unit-length radius in X-Y plane. The center of the domain sits at the origin of the coordinate system. The initial condition for *ϕ*_1_ and *ϕ*_2_ is described by a random function ranging from −0.005 to −0.015 with an average equal to 0.005. The non-zero parameters are set to be: *h* = 0.516, *α* = 8.99 × 10^−3^, *β* = 1 × 10^−2^, *γ* = −*α*, *δ* = 9.1 × 10^−3^, *r*_1_ = 35, *r*_2_ = 0, *k* = 12. From Fig. 2, one can see that for the basic reaction-diffusion model, non-axisymmetric Turing instability occurs, which lead to typical Turing patterns for both components (*ϕ*_1_, *ϕ*_2_). By looking into the third column in Fig. 2, one can see the wave-like oscillation of both *ϕ*_1_ and *ϕ*_2_ along the line X=0.

### Contraction-based colony elasticity

The elasticity model of tissue contraction is described here. The cell colony is modeled as a thin layer of elastic sheet with a thickness *h*_*C*_, and a displacement field **u** within itself, occupying a region of Ω. The plane-stress constitutive law in the elastic sheet can be expressed by^12^:

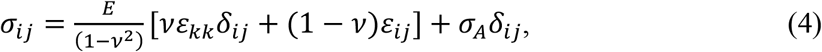

where *E* and *v*, are Young’s modulus, and Poisson’s ratio, respectively. The active contractility of cells is modeled by an active stress *σ*_*A*_*δ*_*ij*_ that resembles a uniform thermal cooling across the cell sheet. Since the amount of the activator and the inhibitor are both low^3^, the stress-free strain caused by them is neglected in Eq. (4). The small deformation strain is written as:

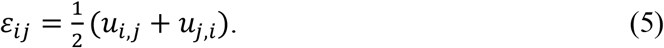

The cell colony interacts with the substrate by exerting traction forces on it through focal adhesions. The traction forces, which in turn are transmitted back into the colony, are balanced with the intercellular stress inside the colony. Hence, the mechanical equilibrium and the boundary condition in this case is given by^21^:

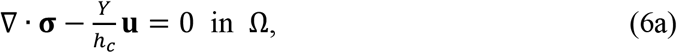

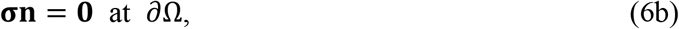

where *Y* is the effective stiffness of the focal adhesion which depends on the stretching strength and the density of the focal adhesions, as well as the stiffness of the substrate. In this article, for simplicity *Y* is assumed to be homogeneous over the whole colony. The two-dimensional tensor **σ**denotes for the Cauchy stress inside the colony, while the vector **n** stands for the outer unit normal of the colony boundary. For the case that *h* ≪ *R*, the traction force −*Y***u** can be averaged along the colony thickness and considered as a body force.

The dimensionless parameters used for the mechanical equilibrium of the cell colony are: *E* = 10^4^, *v* = 0.4, *h*_*C*_ = 0.1, *Y* = 10^4^, *σ*_*A*_ = 500. Solving for the contraction-based elasticity equations (4)–(6) yield the stress field **σ** and the traction stress = *Y***u** that will be used in the following two sections.

### Tension-diffusion coupling: the tension-gradient driven diffusion model

The first proposed mechanism underlying the axisymmetric pattern is a tension-gradient driven diffusion hypothesis. We hypothesize that the diffusion of the activators and the inhibitors within the cell colony is driven by the gradient of “forces” inside, which under this circumstance is the gradient of the intercellular tension −∇*P* = ∇(*σ*_*rr*_ + *σ*_00_)/2. Such gradient maintains the membrane tensions for adjacent cells to help the diffusion of molecules/ions across cell membranes^22^. Thus, the tension-gradient driven diffusion flux is assumed to be proportional to the gradient of the intercellular tension and should exist only when *ϕ*_*i*_ is non-zero:

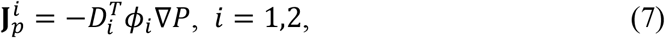

where 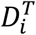 denotes the diffusivity related to the tension-gradient for the *i*-th unknown, which is set to be 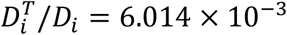. From Eq. (6), one can see that the two signaling molecules are attracted by the increasing of the intercellular tension and repelled by the increasing of the intercellular compression. The total flux is then computed as:

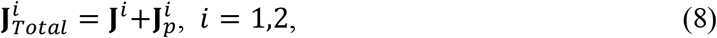

The reaction-diffusion equations and the corresponding zero-flux boundary conditions are then written as:

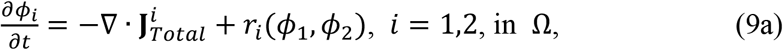

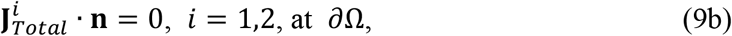

Where *r*_*i*_(*ϕ*_1_, *ϕ*_2_) (*i* = 1,2) are the source terms and are expressed by Eq. (2). The initial conditions are set to be the same with the basic reaction-diffusion model.

Next, we focus on the evolution of *ϕ*_1_ and *ϕ*_2_ driven by the gradient of the intercellular tension. The time evolution of the active level of both signaling molecules are presented in Fig. 3, in which the contours of the active levels of *ϕ*_1_ and *ϕ*_2_ are plotted with respect to several dimensionless time steps. Steered by the axisymmetric distribution of the intercellular tension, the active levels of *ϕ*_1_ and *ϕ*_2_ start from randomness, then gradually evolve into an axisymmetric configuration, and finally arrives at the steady state with alternatively generated peaks and troughs, forming an axisymmetric Turing pattern. At the central region of the colony, *ϕ*_1_ exhibits a high activity, while at the colony periphery, *ϕ*_2_ shows to be more active. Next, we study how the wavenumber-related parameter *k* would affect our results. By setting *k* = 6, 9, 12, we obtain the contours of the steady state solution with respect to different wavenumbers in Fig. 4. From the first column to the last, a clear trend is revealed that a larger *k* will lead to more peaks and troughs to the distribution of the active levels of the two components. When *k* = 12, there are two thorough peaks and troughs; when *k* = 9, we could see one peak with two troughs for *ϕ*_1_ and two peaks with one trough for *ϕ*_2_; when *k* = 6, only one peak and one trough for both *ϕ*_1_ and *ϕ*_2_. At last, we apply the model to predict the activity of the activators and the inhibitors in a square-shaped colony with a dimensionless side length equal to 2. The predicted active levels are plotted in Fig. 5. The results show a 90-degree rotational symmetry of the distribution of the peaks and the troughs, indicating that the tension-gradient driven diffusion is highly regulated by the colony geometry. The effect by the different values of wavenumber-related parameter on the prediction is also fully replicated for the square configuration. However, regardless of the value of *k*, the colony periphery is always the region where *ϕ*_2_ shows its high activity.

**Figure 3.**
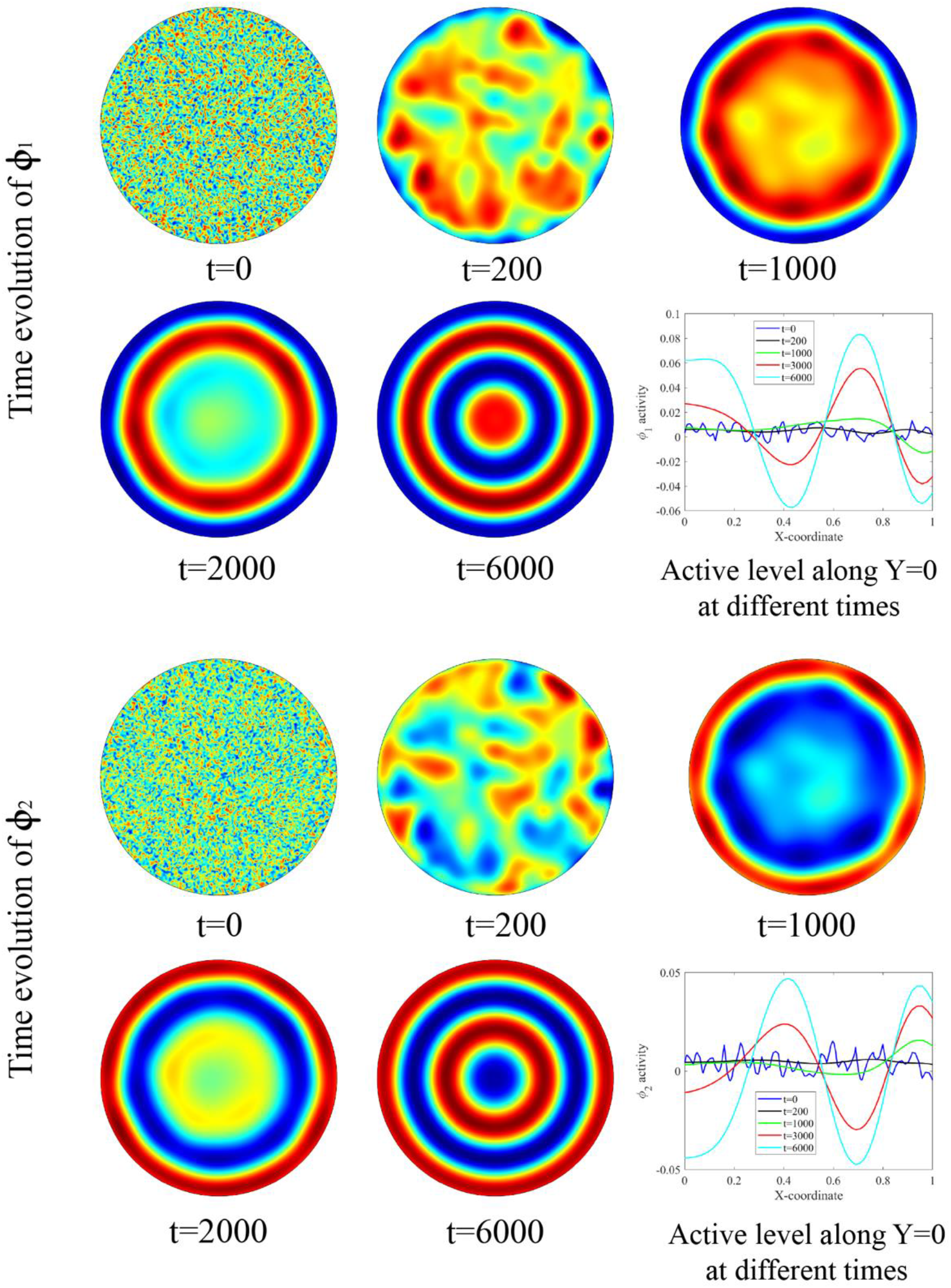
The time evolution contours of the active levels of *ϕ*_1_ and *ϕ*_2_ at different dimensionless time steps are presented (red for large, and blue for small) for a circular colony, by the tension-gradient driven diffusion model. The active levels along X axis for both variables are also plotted.

**Figure 4.**
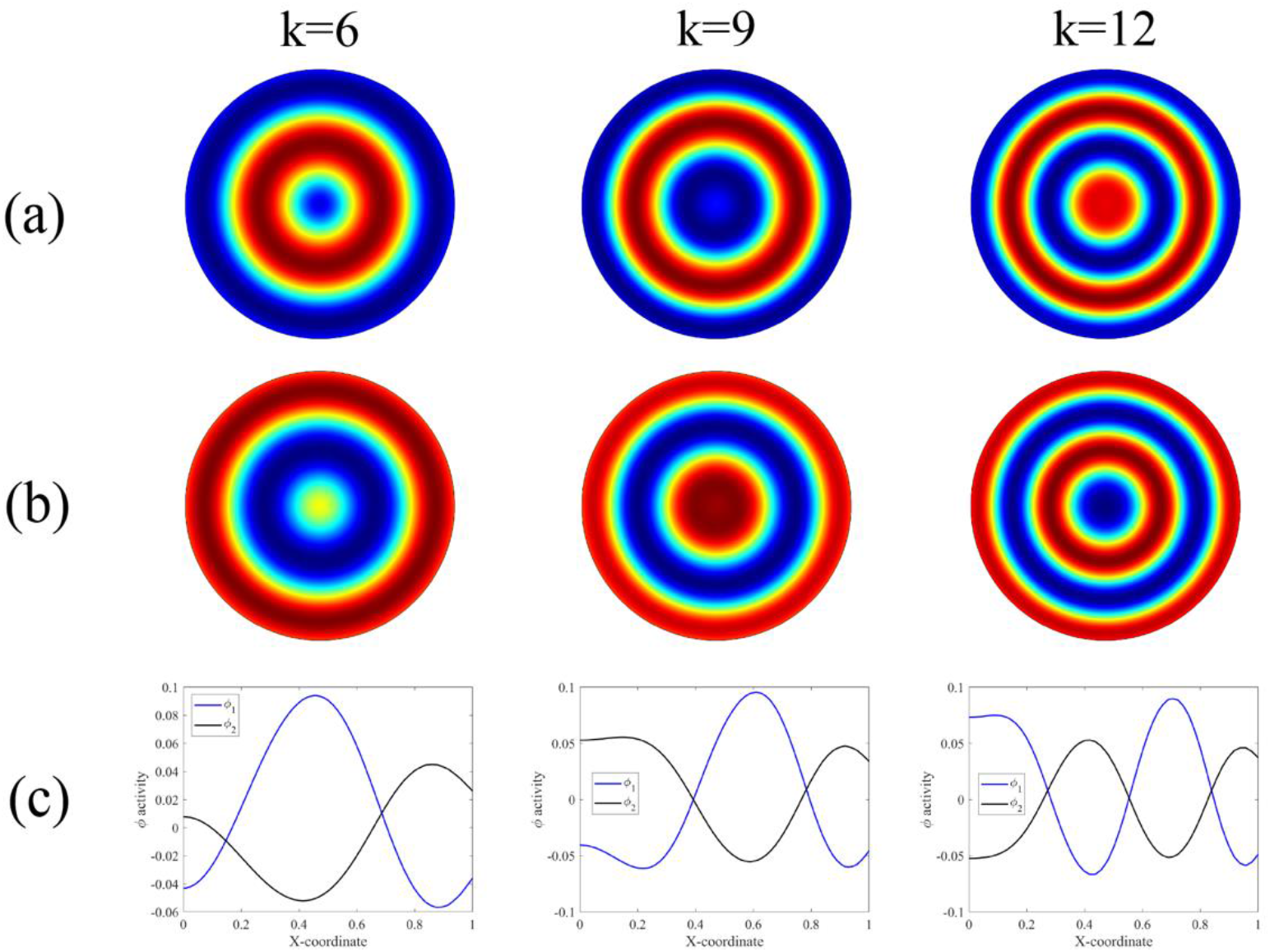
The contours of the steady state solution for the active levels of *ϕ*_1_ and *ϕ*_2_ are plotted for different wavenumbers in a circular-shaped colony. Row (a) is for the results of *ϕ*_1_; Row (b) for *ϕ*_2_; Row (c) shows the distribution of the activity of both *ϕ*_1_ and *ϕ*_2_ along the X-axis.

**Figure 5.**
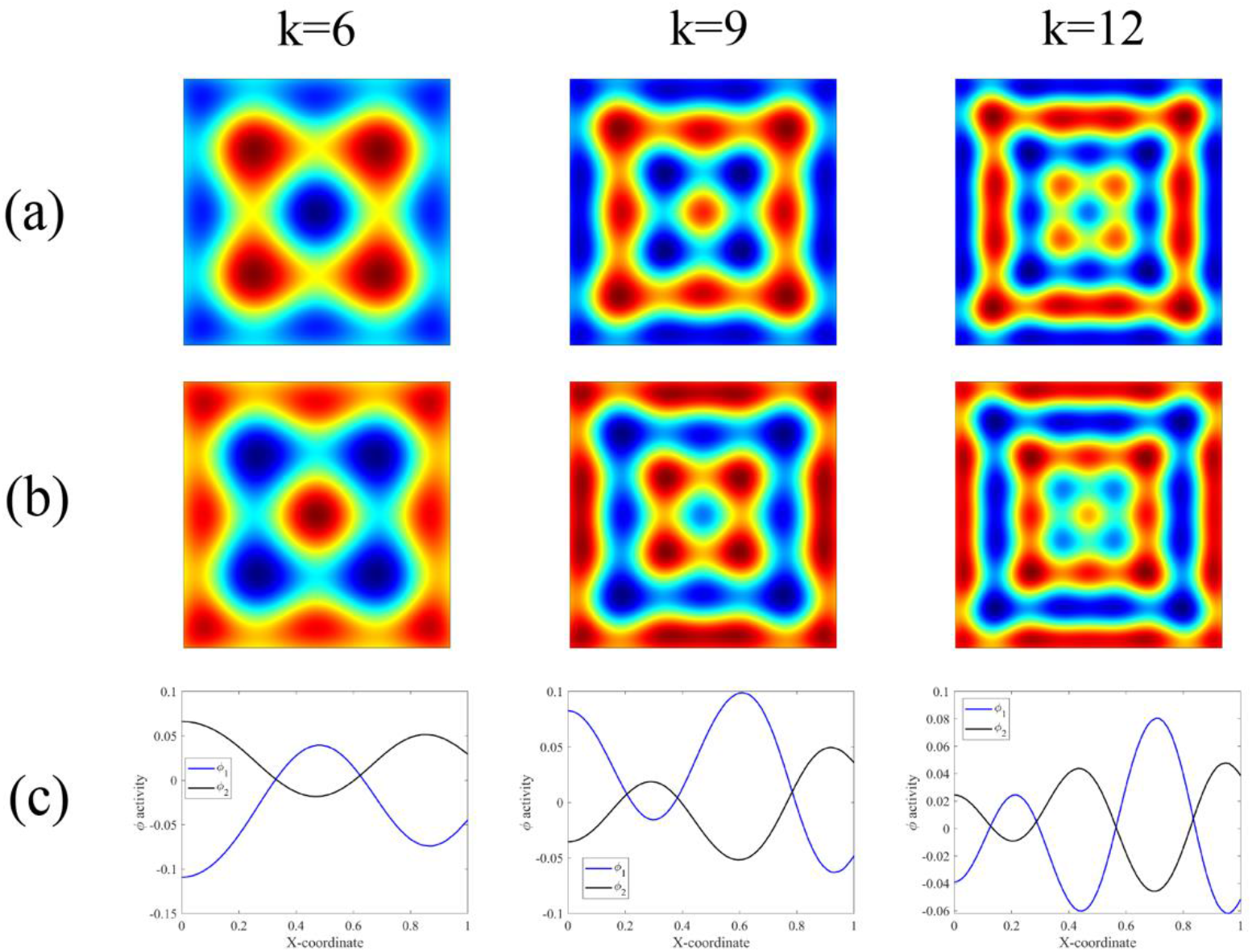
The predicted contours of the steady state solution for the active levels of *ϕ*_1_ and *ϕ*_2_ are plotted for different wavenumbers in a square-shaped colony. Row (a) is for the results of *ϕ*_1_; Row (b) for *ϕ*_2_; Row (c) shows the distribution of the activity of both *ϕ*_1_ and *ϕ*_2_ along the X-axis.

### Traction-force coupling: the traction-force mediated diffusion-reaction model

Unlike the intercellular tension which is developed all over the colony sheet, the traction forces are mainly distributed at the colony periphery that is called boundary layer^12^. The aggregated traction forces in the boundary layer can activate certain signaling pathways that facilitate the generation of the specific signal molecules. By this hypothesis, it is the traction forces, not the intercellular tension that drive the reaction process. Under this hypothesis, the form of the reaction-diffusion equations is still in the form of Eq. (1a) ~(1c), yet some of the parameters are set to be traction-force-dependent. The source term in Eq. (7) is rewritten to be:

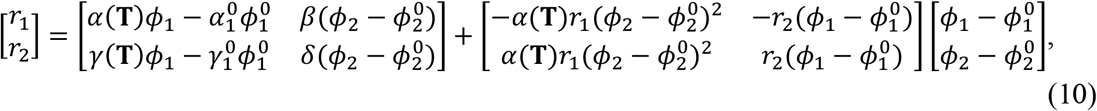

with 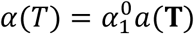, 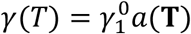, and the traction-force-dependent coefficient *a*(**T**) being assumed as:

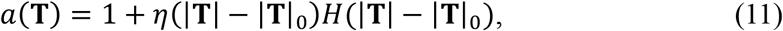

Where *η* = 1/15 is a constant; |**T**|_0_ = 130 acts as a threshold; *H*(·) is the Heaviside function; 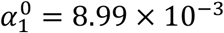, and 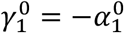; The initial conditions and the rest parameters are again set to be the same with the previous models.

The predictions made by the traction-force mediated diffusion-reaction model are then presented and briefly discussed in this section. The time evolution of the active level of both signaling molecules are presented in Fig. 6. By comparing Fig. 6 with Fig. 3, we see that the distribution of the active levels of *ϕ*_1_ and *ϕ*_2_ predicted by the traction-force mediated model is altered from that predicted by the tension-gradient driven model. Specifically speaking, the location where *ϕ*_1_ shows the high activity and *ϕ*_2_ shows the low activity in the previous model now becomes the place where *ϕ*_2_ shows the high activity and *ϕ*_1_ shows the low activity. This reverse trend is also held even if the value of *k* (see Fig. 7) or the geometry of the colony changes (see Fig. 8, except the last column). Another difference brought by this model is that for the square-shape colony and *k* = 12, the pattern of the predicted active levels changes from rings to strips and dots. This pattern change happens when *k* ≿ 11.5.

**Figure 6.**
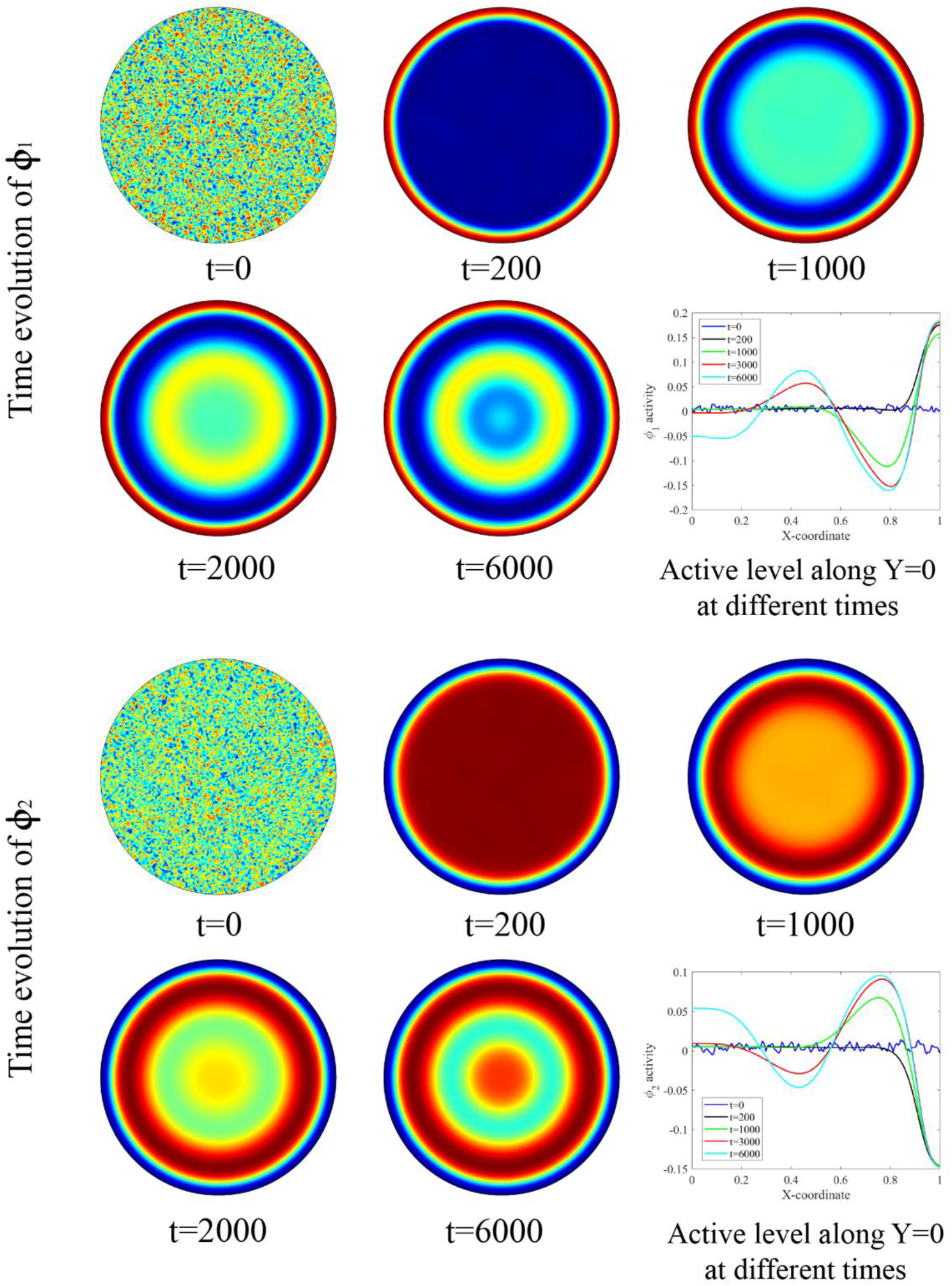
The time evolution contours of the active levels of *ϕ*_1_ and *ϕ*_2_ at different dimensionless time steps are presented (red for large, and blue for small) for a circular colony, by the traction-force mediated diffusion model. The active levels along X axis for both variables are also plotted.

**Figure 7.**
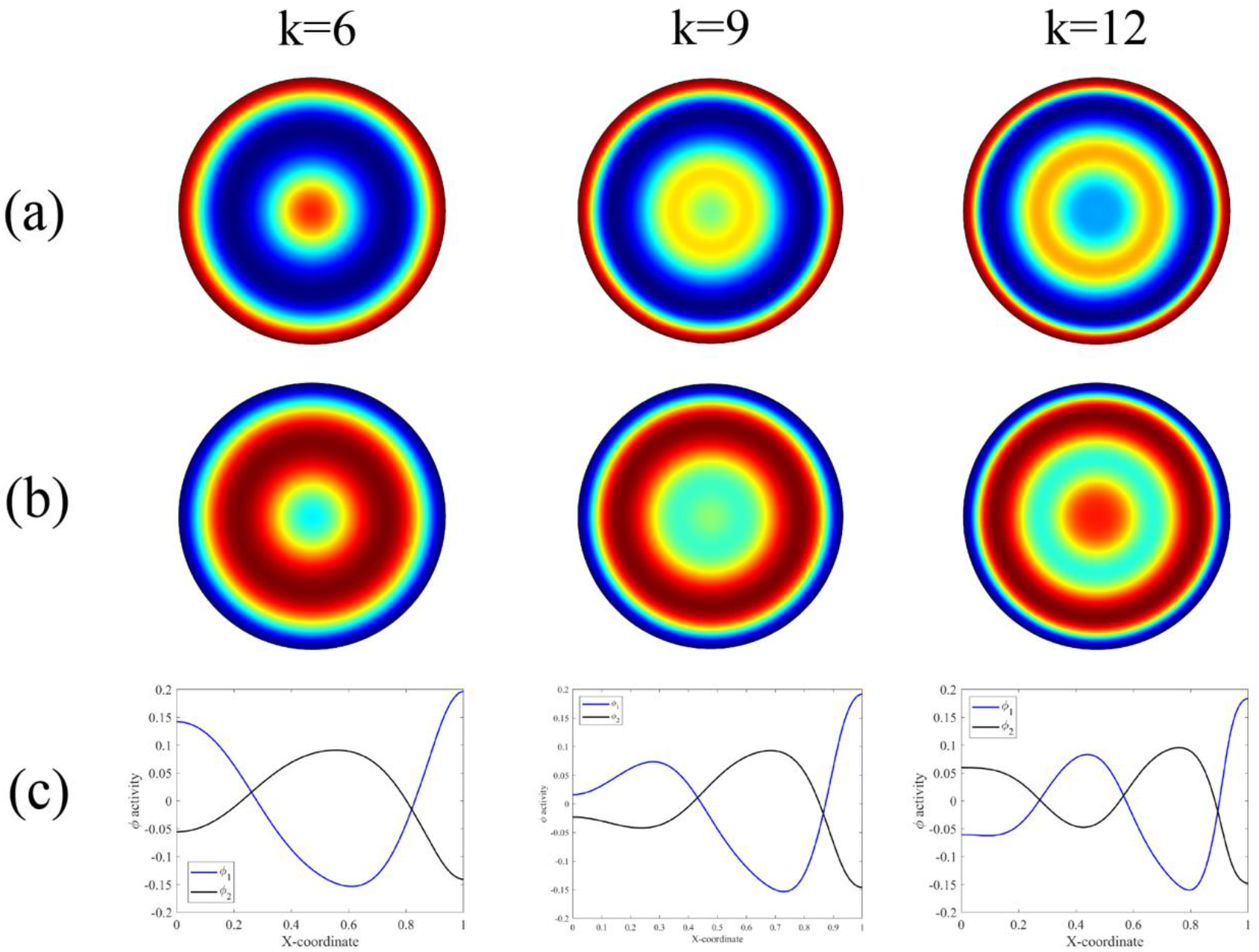
The contours of the steady state solution for the active levels of *ϕ*_1_ and *ϕ*_2_ are plotted for different wavenumbers in a circular-shaped colony. Row (a) is for the results of *ϕ*_1_; Row (b) for *ϕ*_2_; Row (c) shows the distribution of the activity of both *ϕ*_1_ and *ϕ*_2_ along the X-axis.

**Figure 8.**
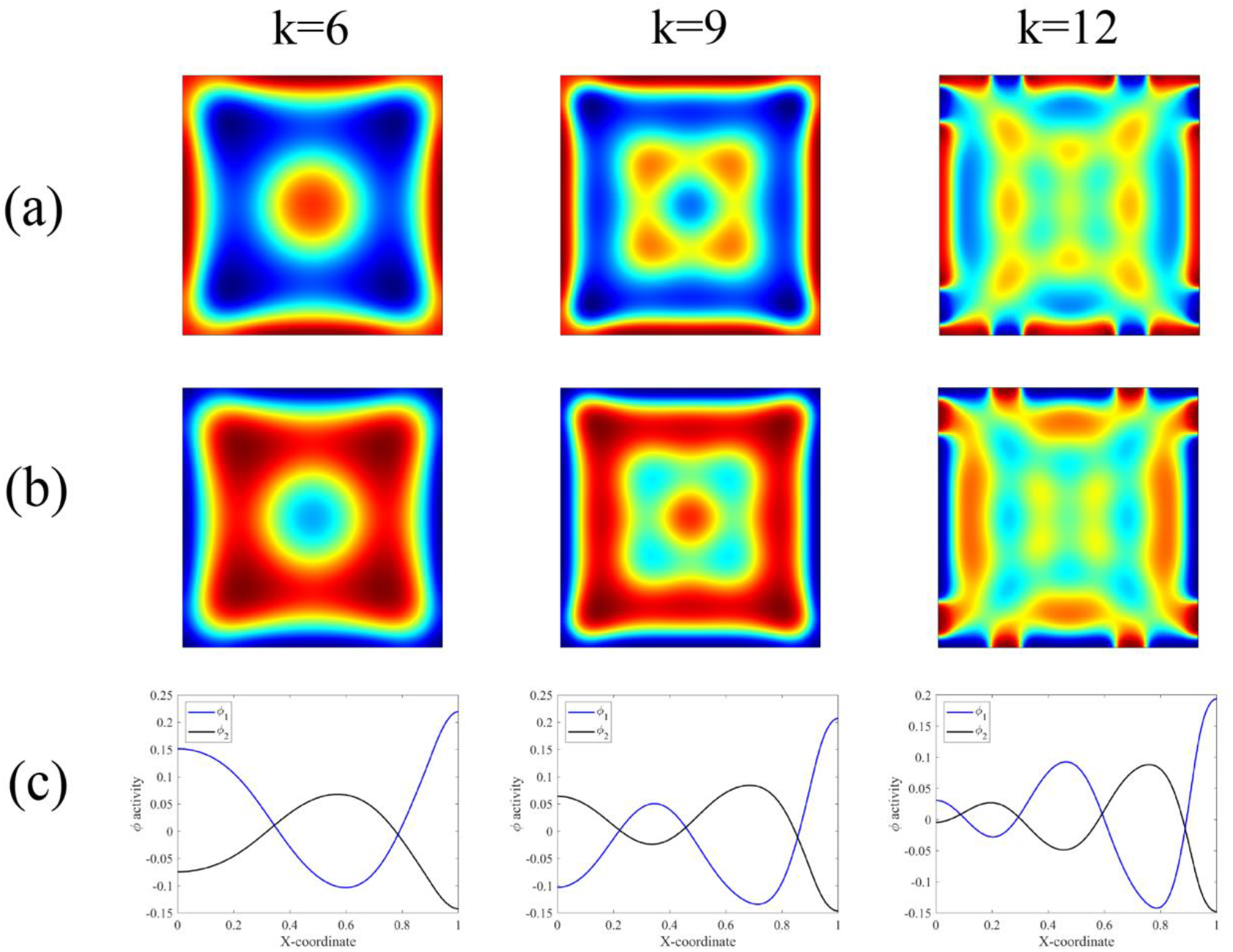
The predicted contours of the steady state solution for the active levels of *ϕ*_1_ and *ϕ*_2_ are plotted for different wavenumbers in a square-shaped colony. Row (a) is for the results of *ϕ*_1_; Row (b) for *ϕ*_2_; Row (c) shows the distribution of the activity of both *ϕ*_1_ and *ϕ*_2_ along the X-axis.

## Conclusion

We raise up two biophysical hypotheses to understand the axisymmetric distribution of the activators and inhibitors in geometrically confined hPSCs-derived microtissues, one is the tension-gradient driven diffusion hypothesis and the other is the traction-force mediated reaction hypothesis. Both theories predict a geometry-regulated distribution of wave-like active levels of the two signaling molecules with random initial conditions. Namely, for the circular-shape colony, an axisymmetric distribution is derived, while a 90-degree rotational symmetric distribution is predicted for the square-shape colony. We note that the locations of the peaks and troughs are altered in the predictions made by the two different theories. The effect of the values of the wavenumber-related parameter is also examined in this article. It is found that a larger value of wavenumber will lead to more waves in the distribution of activity of the two signaling molecules. The authors hope the two biophysical hypotheses can shed some light on the study of mechanisms of early-stage cell fate patterning, particularly using hPSCs-derived in vitro models.

## Acknowledgments

H.Y. acknowledge the funding support from the Southern University of Science and Technology, China (sustech.edu.cn).

## References

(1) Kondo, S.; Miura, T. Reaction-Diffusion Model as a Framework for Understanding Biological Pattern Formation. Science 2010, 329 (5999), 1616. https://doi.org/10.1126/science.1179047.

(2) Warmflash, A.; Sorre, B.; Etoc, F.; Siggia, E. D.; Brivanlou, A. H. A Method to Recapitulate Early Embryonic Spatial Patterning in Human Embryonic Stem Cells. Nat. Methods 2014, 11 (8), 847–854. https://doi.org/10.1038/nmeth.3016.

(3) Tewary, M.; Ostblom, J.; Prochazka, L.; Zulueta-Coarasa, T.; Shakiba, N.; Fernandez-Gonzalez, R.; Zandstra, P. W. A Stepwise Model of Reaction-Diffusion and Positional Information Governs Self-Organized Human Peri-Gastrulation-like Patterning. Development 2017, 144 (23), 4298–4312. https://doi.org/10.1242/dev.149658.

(4) Xue, X.; Sun, Y.; Resto-Irizarry, A. M.; Yuan, Y.; Aw Yong, K. M.; Zheng, Y.; Weng, S.; Shao, Y.; Chai, Y.; Studer, L.; Fu, J. Mechanics-Guided Embryonic Patterning of Neuroectoderm Tissue from Human Pluripotent Stem Cells. Nat. Mater. 2018, 17 (7), 633–641. https://doi.org/10.1038/s41563-018-0082-9.

(5) Heemskerk, I.; Burt, K.; Miller, M.; Chhabra, S.; Guerra, M. C.; Liu, L.; Warmflash, A. Rapid Changes in Morphogen Concentration Control Self-Organized Patterning in Human Embryonic Stem Cells. eLife 2019, 8, e40526. https://doi.org/10.7554/eLife.40526.

(6) Xie, T.; Kang, J.; Pak, C.; Yuan, H.; Sun, Y. Temporal Modulations of NODAL, BMP, and WNT Signals Guide the Spatial Patterning in Self-Organized Human Ectoderm Tissues. Matter 2020. https://doi.org/10.1016/j.matt.2020.04.012.

(7) Haremaki, T.; Metzger, J. J.; Rito, T.; Ozair, M. Z.; Etoc, F.; Brivanlou, A. H. Self-Organizing Neuruloids Model Developmental Aspects of Huntington’s Disease in the Ectodermal Compartment. Nat. Biotechnol. 2019, 37 (10), 1198–1208. https://doi.org/10.1038/s41587-019-0237-5.

(8) Sanderson, A. R.; Kirby, R. M.; Johnson, C. R.; Yang, L. Advanced Reaction-Diffusion Models for Texture Synthesis. J. Graph. Tools 2006, 11 (3), 47–71. https://doi.org/10.1080/2151237X.2006.10129222.

(9) Zhang, W.; Wang, S.; Lin, M.; Han, Y.; Zhao, G.; Lu, T. J.; Xu, F. Advances in Experimental Approaches for Investigating Cell Aggregate Mechanics. Acta Mech. Solida Sin. 2012, 25 (5), 473–482. https://doi.org/10.1016/S0894-9166(12)60042-1.

(10) Ji, B.; Bao, G. Cell and Molecular Biomechanics: Perspectives and Challenges. Acta Mech. Solida Sin. 2011, 24 (1), 27–51. https://doi.org/10.1016/S0894-9166(11)60008-6.

(11) Li, B.; Sun, S. X. Coherent Motions in Confluent Cell Monolayer Sheets. Biophys. J. 2014, 107 (7), 1532–1541. https://doi.org/10.1016/j.bpj.2014.08.006.

(12) Zhao, T.; Zhang, Y.; Wei, Q.; Shi, X.; Zhao, P.; Chen, L.-Q.; Zhang, S. Active Cell-Matrix Coupling Regulates Cellular Force Landscapes of Cohesive Epithelial Monolayers. Npj Comput. Mater. 2018, 4 (1), 10. https://doi.org/10.1038/s41524-018-0069-8.

(13) Zhang, Y.; Shi, X.; Zhao, T.; Huang, C.; Wei, Q.; Tang, X.; Santy, L. C.; Saif, M. T. A.; Zhang, S. A Traction Force Threshold Signifies Metastatic Phenotypic Change in Multicellular Epithelia. Soft Matter 2019, 15 (36), 7203–7210. https://doi.org/10.1039/C9SM00733D.

(14) Yuan, H.; Marzban, B.; Parker, K. K. Myofibrils in Cardiomyocytes Tend to Assemble Along the Maximal Principle Stress Directions. J. Biomech. Eng. 2017, 139 (12). https://doi.org/10.1115/1.4037795.

(15) Marzban, B.; Kang, J.; Li, N.; Sun, Y.; Yuan, H. A Contraction–Reaction–Diffusion Model: Integrating Biomechanics and Biochemistry in Cell Migration. Extreme Mech. Lett. 2019, 32, 100566. https://doi.org/10.1016/j.eml.2019.100566.

(16) Marzban, B.; Yi, X.; Yuan, H. A Minimal Mechanics Model for Mechanosensing of Substrate Rigidity Gradient in Durotaxis. Biomech. Model. Mechanobiol. 2018, 17 (3), 915–922. https://doi.org/10.1007/s10237-018-1001-3.

(17) Nelson, C. M.; Jean, R. P.; Tan, J. L.; Liu, W. F.; Sniadecki, N. J.; Spector, A. A.; Chen, C. S. Emergent Patterns of Growth Controlled by Multicellular Form and Mechanics. Proc. Natl. Acad. Sci. 2005, 102 (33), 11594–11599. https://doi.org/10.1073/pnas.0502575102.

(18) Miura, T.; Shiota, K.; Morriss-Kay, G.; Maini, P. K. Mixed-Mode Pattern in Doublefoot Mutant Mouse Limb—Turing Reaction–Diffusion Model on a Growing Domain during Limb Development. J. Theor. Biol. 2006, 240 (4), 562–573. https://doi.org/10.1016/j.jtbi.2005.10.016.

(19) Onimaru, K.; Marcon, L.; Musy, M.; Tanaka, M.; Sharpe, J. The Fin-to-Limb Transition as the Re-Organization of a Turing Pattern. Nat. Commun. 2016, 7 (1), 11582. https://doi.org/10.1038/ncomms11582.

(20) Barrio, R. A Two-Dimensional Numerical Study of Spatial Pattern Formation in Interacting Turing Systems. Bull. Math. Biol. 1999, 61 (3), 483–505. https://doi.org/10.1006/bulm.1998.0093.

(21) Mertz, A. F.; Banerjee, S.; Che, Y.; German, G. K.; Xu, Y.; Hyland, C.; Marchetti, M. C.; Horsley, V.; Dufresne, E. R. Scaling of Traction Forces with the Size of Cohesive Cell Colonies. Phys. Rev. Lett. 2012, 108 (19), 198101. https://doi.org/10.1103/PhysRevLett.108.198101.

(22) Yang, Y. Durotaxis Index of 3T3 Fibroblast Cells Scales with Stiff-to-Soft Membrane Tension Polarity. 12.

